# Boolean Logic-responsive FRET Biosensors via Genetically Encoded Autonomous Compilation

**DOI:** 10.64898/2026.09.24.753966

**Authors:** Hyerim Son, Jack W. Hoye, Michael Khawand, Murial L. Ross, Ryan Gharios, Cole A. DeForest

## Abstract

Förster resonance energy transfer (FRET) is commonly used to monitor protein-protein interactions *in situ*. The high spatiotemporal resolution and facile implementation inside complex molecular environments have spearheaded FRET’s widespread adoption in biosensing. Despite these advantages, current FRET biosensors are largely restricted to the detection of the presence/absence of *individual* inputs and are thus unable to sense several multiplexable inputs simultaneously within complex milieu of biological environments. In this work, we introduce a generalizable strategy to construct genetically encoded protein-based FRET biosensors capable of recognizing *multiple* inputs following Boolean logic-type (YES/OR/AND) operations. These topologically specified FRET sensors powerfully expand the input capacity in sensing protein-protein interactions while providing a user-programmable platform for monitoring heterogeneous biological activities both *in vitro* and in living cells.

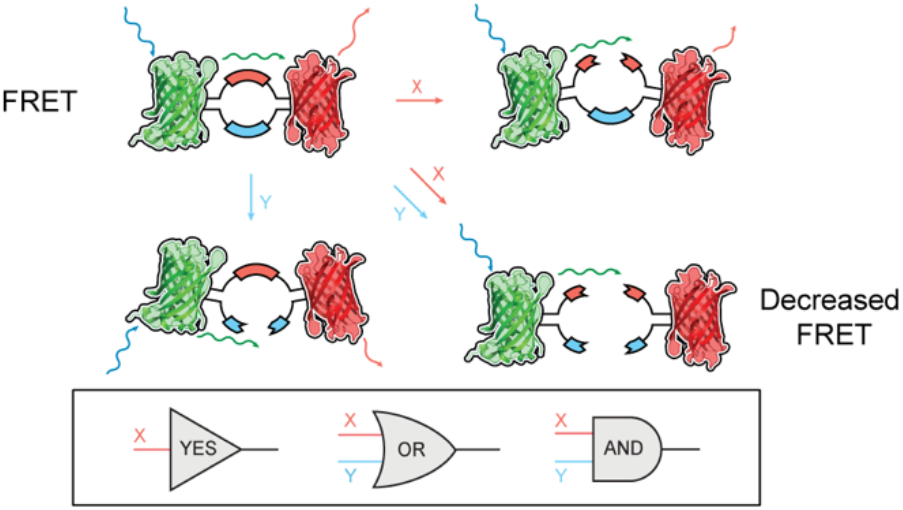

## Main Text

Biosensors convert a molecular recognition event into a measurable output, and their underlying (bio)chemistry determines what they report and where they can operate^1,2^. Nucleic acid circuits^3^, whole-cell bacterial sensors^4^, and protein-based detectors^5^ have each been developed for analyte detection, diagnostics, and the readout of activity inside engineered cells. Among these, Förster resonance energy transfer (FRET) biosensors quantify species proximity through a physical relationship well suited to intracellular measurement^6^. Energy transfer from donor to acceptor scales as *r*^*−6*^, where *r* is the distance separating the pair, such that nanometer-scale changes in separation yield large changes in FRET signal^7^ that can be detected within living systems at subcellular resolution^8–10^. Genetically encoded FRET biosensors based on fluorescent proteins are particularly enabling, in that they can be continuously synthesized within the exact biological context for which they will be employed^11–14^. Though such systems have permitted quantification of protease activity in tumors^15,16^, coenzyme homeostasis^17^, and metabolite flux^18,19^, virtually all FRET biosensors reported to date determine the presence or absence of an *individual* cue, as their donor and acceptor are joined by a single responsive element. As cellular decisions are rarely made based on a single input, systems capable of sensing *multiple* inputs simultaneously within biological environments remain of great interest^20^.

In this manuscript, we introduce a generalizable strategy to genetically construct protein-based FRET biosensors that respond to user-programmable combinations of biological inputs via Boolean logic (YES/OR/AND). We have previously exploited controlled molecular topology to program Boolean-based biomaterial degradation^21,22^ and cargo delivery^23,24^, though such methods have not yet been applied in biosensors. Here, we co-express two fluorophores of interest – EGFP as the donor and mScarlet3 as the acceptor – connected by a topologically specified degradable linker sensitive to distinct protein inputs. These biosensors are “autonomously compiled” from a single open reading frame via spontaneous intramolecular ligations following expression^25^. When the linker region is intact, the fluorescent proteins are held in close proximity, permitting efficient FRET transfer to occur; when cleaved, FRET decreases substantially (**Fig. 1a**). The two inputs used throughout are an evolved Sortase [eSrtA(2A9)] transpeptidase (which recognizes the polypeptide motif LAET*↓G*) and a potyviral tobacco etch virus (TEV) protease (which recognizes ENFLYQ*↓S*) (**Fig. 1b**). As the logical operation follows from connectivity rather than from the chemical identity of the motifs, arranging these two sites within the linker yields the three architectures examined here: a single scissile site (YES), two sites connected in series (OR, logic symbol ∨), and two sites connected in parallel (AND, logic symbol ∧) (**Fig. 1c-e**). Each biosensor was first expressed recombinantly in *E. coli* and characterized in solution, then stably expressed and validated in mammalian cells.

**Figure 1.**
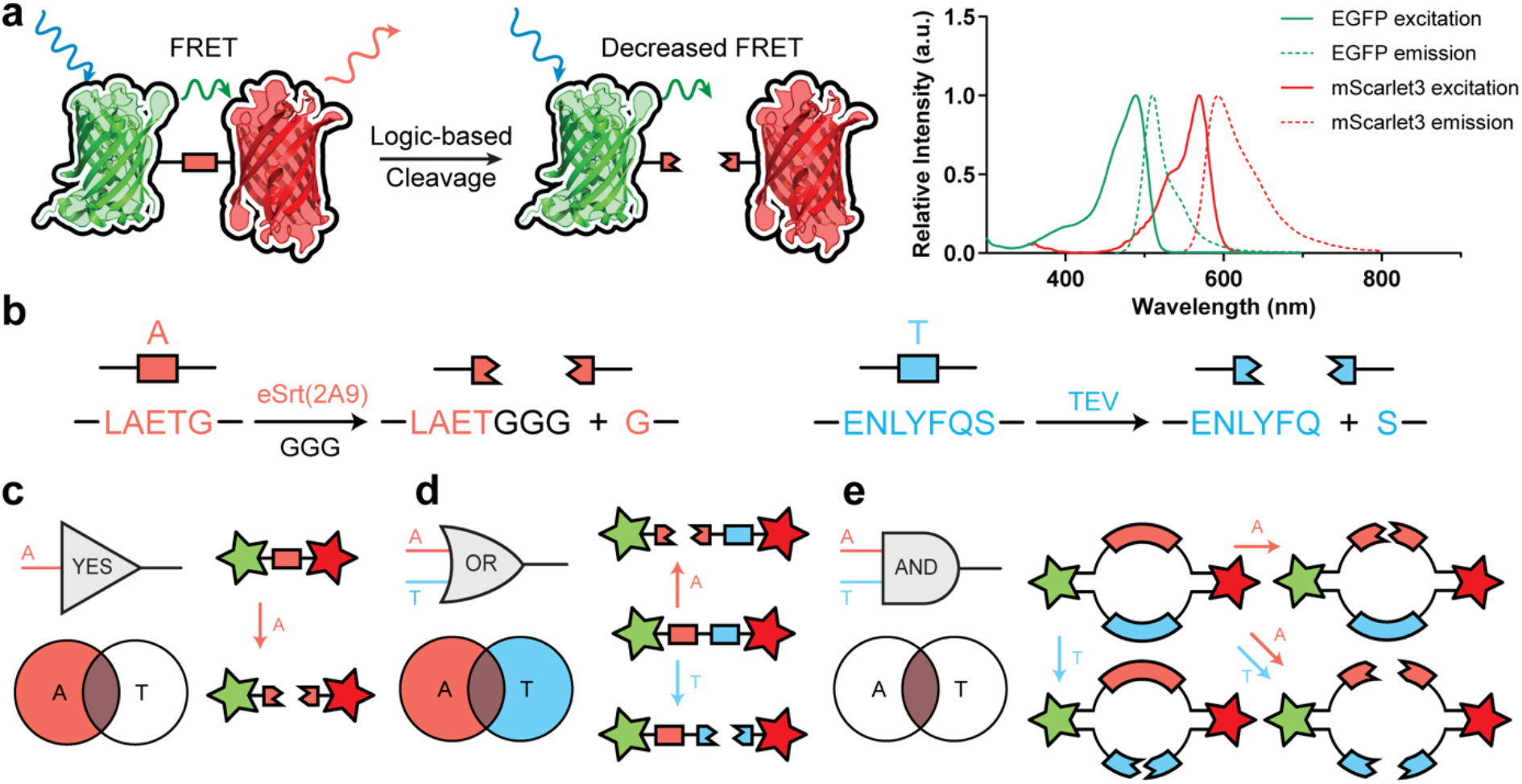
Autonomously compiled Boolean logic-gated FRET protease sensors. (a) (left) Scheme of FRET ON/OFF mechanism after gated scission, (right) excitation/emission spectra of FRET donor (EGFP) and acceptor (mScarlet3) used in these studies. (b) eSrtA(2A9) and TEV protease inputs and their substrates motifs used in these studies. (c-e) Boolean logic-gated linkers between EGFP (green star) and mScarlet3 (red star): (c) a YES-gate with a single protease-recognizing site, (d) an OR gate (logic symbol ∨) with two protease-labile moieties linked in series, and (e) an AND-gate (logic symbol ∧) with two protease substrates connected in parallel. (Adapted from Gharios et al., *Nat. Chem. Bio*., **2025**.^25^ Copyright 2025 Nature. Reproduced with permission).

We focused our initial efforts on the construction of the simplest logical operation: a YES-gated system responsive to eSrtA(2A9) (construct denoted as A). Here, EGFP and mScarlet3 were genetically linked as a fusion pair via a flexible linker containing the LAETG motif (**Fig. 2a**). The 6xHis-tagged protein was expressed in *E. coli* and purified via immobilized metal affinity chromatography (IMAC) and size-exclusion chromatography (SEC) (**Fig. S1-S3**). After treatment with all available protease combinations [i.e., eSrtA(2A9) alone denoted as _A_, TEV alone denoted as _T_, eSrtA(2A9) and TEV in tandem denoted as _AT_, and no treatment], samples were analyzed by sodium dodecyl sulfate-polyacrylamide gel electrophoresis (SDS-PAGE) (**Fig. 2b**). As expected, samples treated with eSrtA(2A9) (i.e., _A_ and _AT_ groups) were cleaved, indicated by the robust mass shift and the presence of two distinct protein products. These findings also indicate that crosstalk between the TEV and eSrtA(2A9) are undetectable, asserting these input cues as orthogonal to one another. To confirm these results further, input-treated products were subjected to liquid chromatography-tandem mass spectrometry (LC-MS); results show the cleavage follows the expected pathway for canonical eSrtA(2A9)-mediated transpeptidation (**Fig. 2c**).

**Figure 2.**
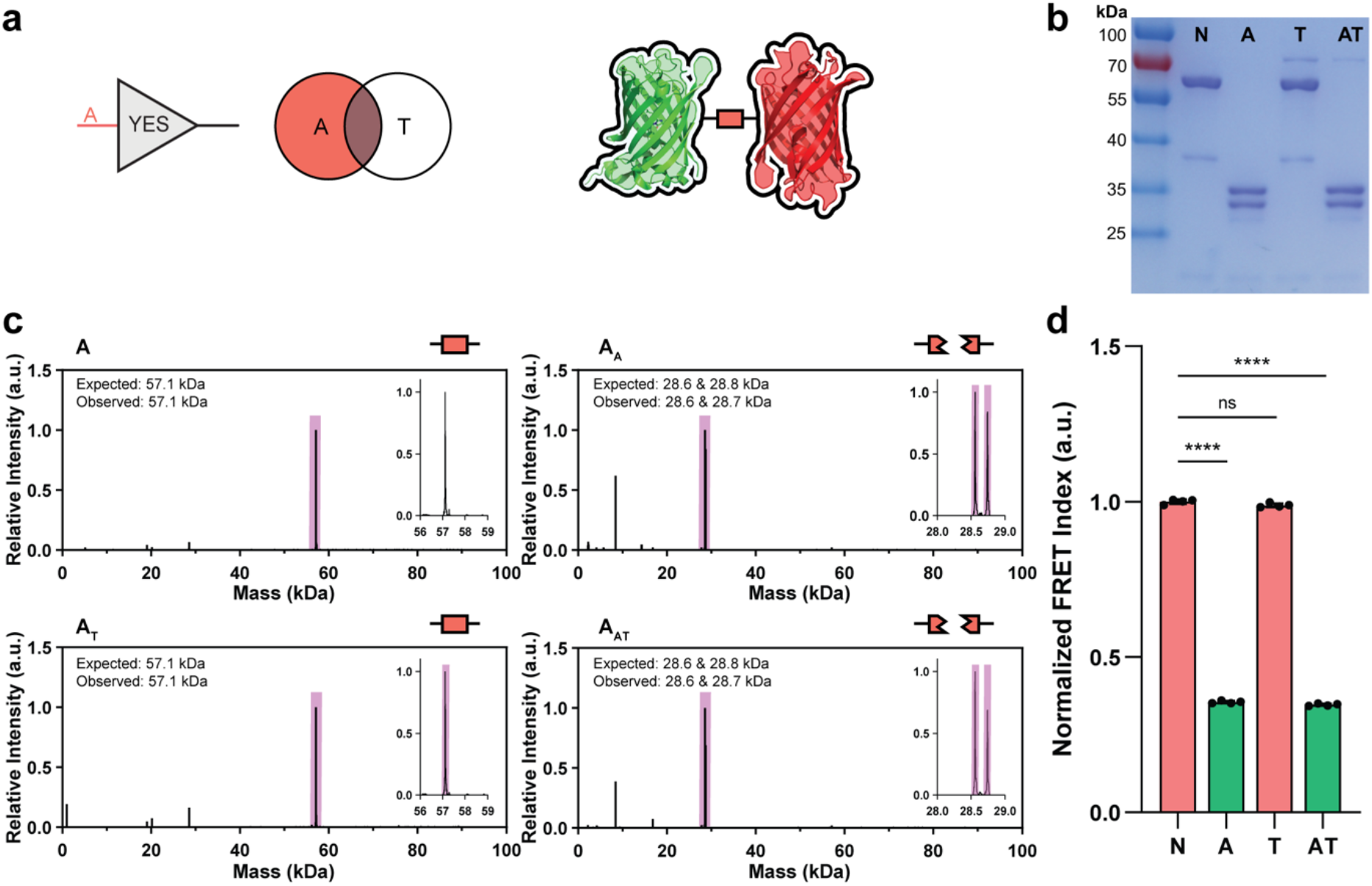
YES-gated FRET sensor response. (a) The A biosensor is cleaved in response to eSrtA(2A9), as confirmed by (b) SDS-PAGE and (c) LC-MS analysis. (d) Normalized FRET index following protease inputs. _N_ indicates no treatment, _A_ indicates eSrtA(2A9), and _T_ indicates TEV. Bars for conditions in which biosensor response are expected are colored green, while those unexpected are colored red. One-way ANOVA test with Šídák’s multiple comparisons test comparing each condition to the control, **** = p<0.0001. Unless marked, all other conditions are not significant.

To assay the functional response of the YES-gated system, variable protease-treated samples were assayed for FRET using the excitation wavelength of EGFP (∼480 nm) while monitoring emission for both EGFP (∼510 nm) and mScarlet3 (∼590 nm). The FRET Index ratio was calculated as:

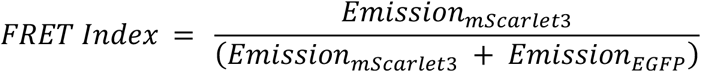

*w*ith the Normalized FRET Index reported relative to the highest FRET Index within the group. Compared to the non-treated control, constructs exposed to TEV showed negligible reduction in FRET activity. Conversely, FRET activity dropped significantly with _A_ and _AT_ treatments, with the mean FRET ratio achieving roughly a third of the non-treated group (**Fig. 2d**).

Building on these successes, we next synthesized an OR-gated construct responsive to either eSrtA(2A9) or TEV inputs (denoted A∨*T*), assembled with a single linker between the FRET pairs bearing both protease recognition domains connected in series (**Fig. 3a, S4-S6**). As designed, the OR gate showed strong responsiveness to treatment with either protease, as analyzed via SDS-PAGE gel shift assay (**Fig. 3b**). LC-MS confirms expected scission occurs with the _A, T_, and _AT_ treatments, as well as the expected mass loss from the intervening linker region diffusing away upon treatment with both proteases simultaneously (**Fig. 3c**). FRET response also adhered to expectations and mirrored the results of the YES gate. All treatment conditions yielded a near-identical drop in the FRET ratio such that the final FRET ratio represented 38 ± 2% of the untreated condition (**Fig. 3d**).

**Figure 3.**
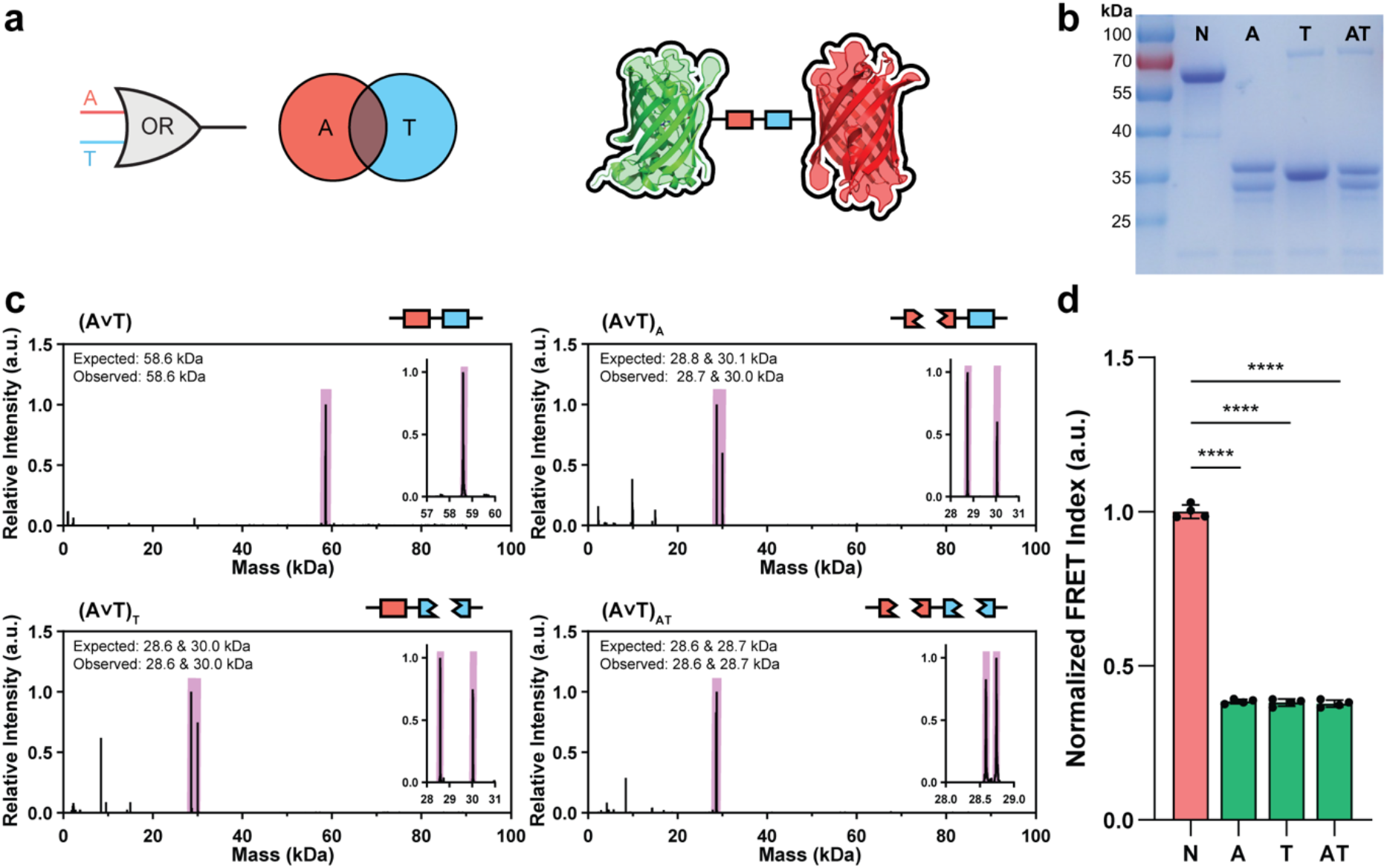
OR-gated FRET sensor response. (a) The A∨*T* biosensor is cleaved in response to TEV OR eSrtA(2A9), as confirmed by (b) SDS-PAGE and (c) LC-MS analysis. (d) Normalized FRET index following protease inputs. _N_ indicates no treatment, _A_ indicates eSrtA(2A9), and _T_ indicates TEV. Statistics, histogram coloring, and treatment abbreviations are the same as in Figure 2.

To achieve AND-gated computation, fluorescent proteins within the A∧*T* protein construct are linked with scissile motifs connected in parallel, achieved here via N-to-C cyclization (**Fig. 4a, S7-S9**). This recursive linker design allows each motif to be degraded separately and orthogonally to one another; individual cleavage of either motif linearizes the initially cyclic species, while complete separation requires degradation of both. Protein cyclization was achieved via the “split-intein circular ligation of peptides and proteins” (SICLOPPS^26^) method utilizing the Cfa system^27^; here, Cfa fragments appended in reversed orientation (i.e., Cfa^C^ at the N terminus of the FRET species, Cfa^N^ at its C terminus) reassociate into an active intein that ligates the construct’s own termini, rapidly and near-scarlessly cyclizing the intervening protein while excising itself from the primary sequence. This autocatalytic topological assembly process is highly efficient in both *E. coli* and mammalian cells. After bacterial expression and IMAC/SEC-based purification, the A∧*T* species was subjected to SDS-PAGE analysis (**Fig. 4b**). Here, samples treated with either the _A_ or the _T_ input remained intact, though their apparent molecular weight shifted slightly, consistent with their transition from a cyclic to linear construct. When exposed to both the eSrtA(2A9) and TEV proteases, EGFP and mScarlet3 were proteolytically separated into distinct species, observed by the appearance of lower molecular weight bands (∼35 kDa observed). LC-MS analysis indicates the expected masses corresponding to the intact and cyclic FRET species in the absence of treatment, the linearized constructs upon either _A_ or _T_ treatment, and then fully disassociated fluorescent proteins upon treatment with both inputs (**Fig. 4c**).

**Figure 4.**
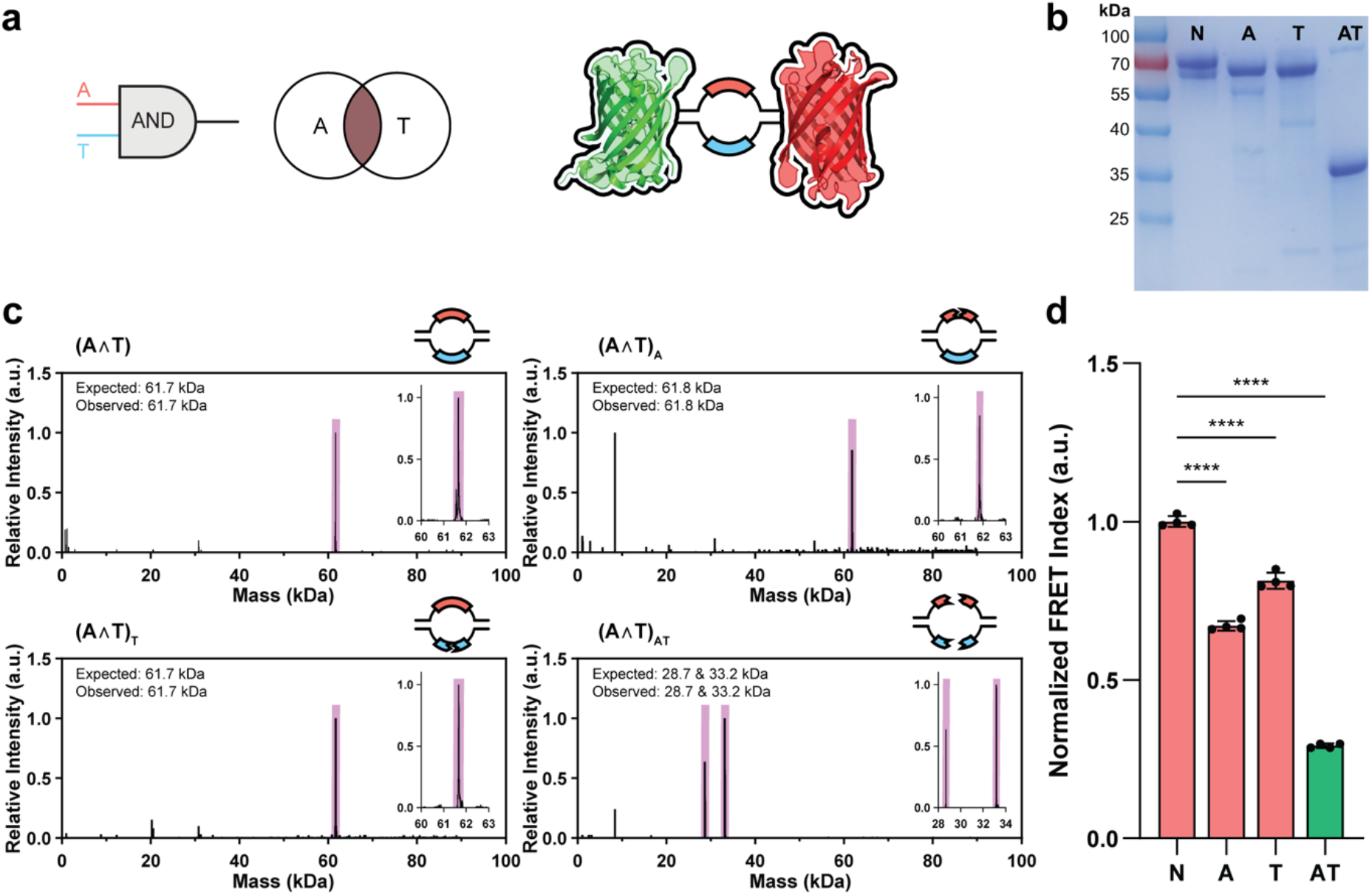
AND-gated FRET sensor response. (a) The A∧*T* biosensor is fully cleaved in response to TEV AND eSrtA(2A9), as confirmed by (b) SDS-PAGE and (c) LC-MS analysis. (d) Normalized FRET index following protease inputs. _N_ indicates no treatment, _A_ indicates eSrtA(2A9), and _T_ indicates TEV. Statistics, histogram coloring, and treatment abbreviations are the same as in Figure 2.

The cyclic structure of the A∧*T* gate is also highly constrained and limits the conformational states available to the protein structure, which has detectable ramifications on the FRET response. The cyclic species’ FRET ratio is higher than that of either of the linear gates due to this constrained conformation, which restricts the fluorophore separation and notably improves the FRET efficiency. Upon linearization in response to either protease cue delivered in isolation, the FRET ratio noticeably decreases due to relaxation of the gate’s conformation (**Fig. 4d**). Differences in linker length are also detectable using the FRET ratio, where the retained linker of the A treatment is longer than that of the T, causing a lower FRET ratio to be observed. When treated with both cues, the FRET ratio decays to a similar baseline as the OR and YES gates, consistent with LC-MS results showing total dissociation of the fluorophores from one another.

We next asked whether the same constructs would compile and compute inside living cells, where the fluorophores, proteases, and enzymatic cofactors are all supplied by the host. HEK293T cells were stably modified to individually express all gate combinations (i.e., A, A∨*T*, or A∧*T*) in a doxycycline-inducible manner (**Fig. 5a**). To faithfully replicate the results shown in solution, DNA encoding an identical amino acid sequence for each FRET construct was incorporated into the host cell genome via Sleeping Beauty transposition^28^. After 24 hours of doxycycline treatment, confocal imaging confirmed biosensor expression, with emission in both channels and a high FRET ratio prior to protease delivery. As donor and acceptor are translated from a single open reading frame, the two fluorophores are necessarily present in equimolar amounts.

**Figure 5.**
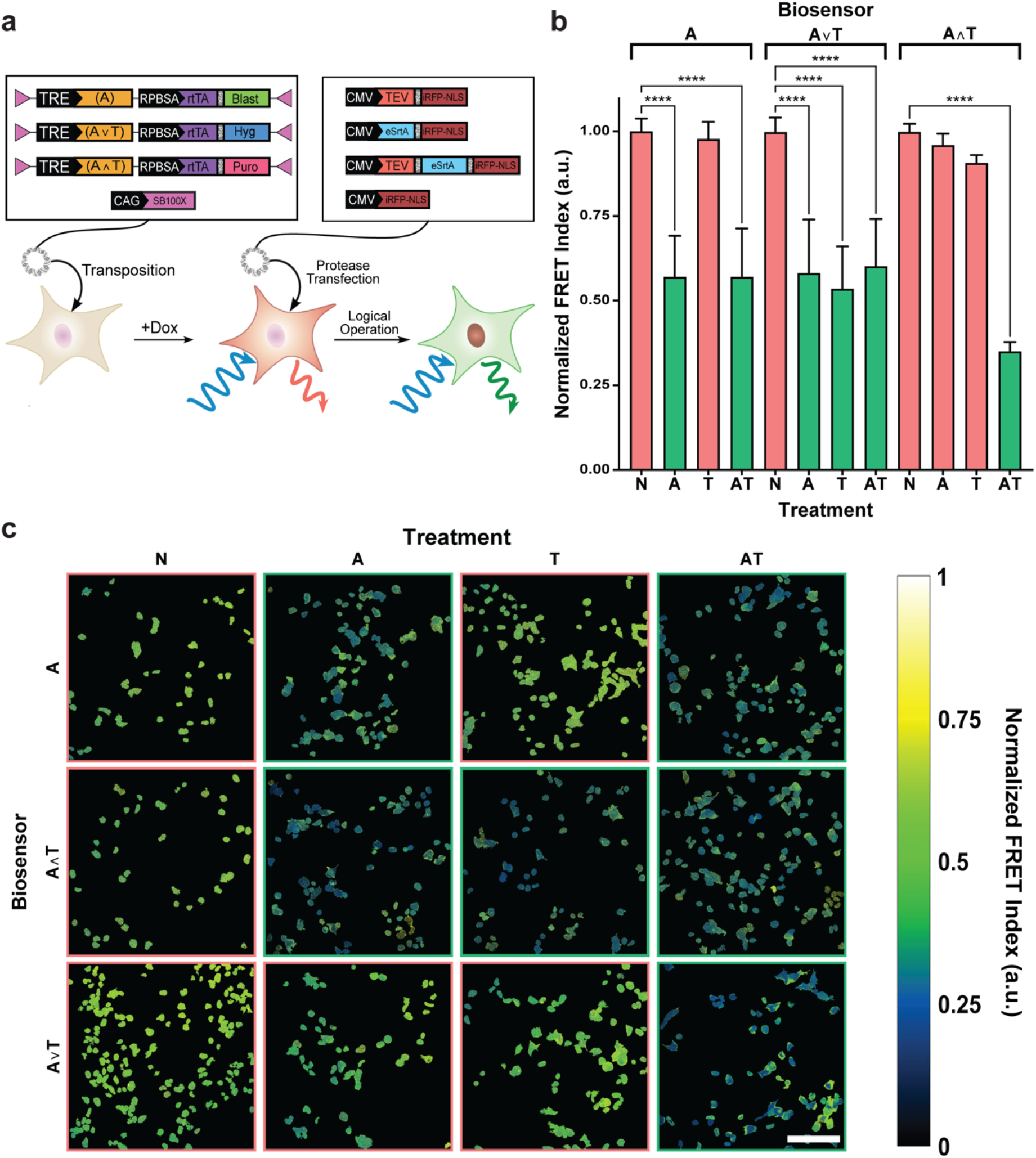
Mammalian logic-gated FRET biosensors *in cellulo*. (a) Overview of gene design and experimental workflow. Sleeping Beauty transposons are first integrated into HEK393T cells via transposition, SB-ITR sequences are directionally annotated with a pink triangle. After logic-gated biosensors are integrated and cells are treated with doxycycline, all cells display a high FRET index. After transient transfection with protease-encoding plasmids, FRET ratio declines in the expected manner programmed by gate architecture. (b) Normalized FRET ratio of well-transfected cells, analyzed via flow cytometry. Statistics, histogram coloring, and treatment abbreviations are the same as in Figure 2. (c) Representative images of biosensor-encoding cell nuclei following protease expression. Images where FRET ratio is expected to decline are outlined in green while those who are expected to remain constant are outlined in red. Color bar indicates intensity values of the calculated FRET ratio at any given point. Scale bar = 100 µm.

Following gate expression, cells were transfected with plasmids encoding (i) no protease, (ii) eSrtA(2A9), (iii) TEV, or (iv) both protease inputs, each co-expressed with a spectrally distinct reporter protein – a near-infrared fluorescent protein (iRFP) genetically fused to a nuclear localization signal – to monitor transgene expression levels within individual cells. Following transfection and quantification by flow cytometry and confocal microscopy, all gate architectures faithfully preserved and mirrored the results seen in solution (**Fig. 5b-c**). Interestingly, proteases expressed inside of living cells did not perform equally well to those isolated in solution with respect to facilitating linker degradation. While TEV expression in cells led to robust scission of its target sequence at virtually all detectable expression levels, eSrtA(2A9) required a much greater degree of overexpression to facilitate cleavage of its target sequence. TEV requires a reducing environment to function^29,30^, a condition the mammalian cytosol natively maintains^31^, so its activity is not limited in this setting, consistent with the robust scission we observe across nearly all detectable expression levels. Contrastingly, eSrtA(2A9) requires a Ca^2+^ cofactor and an N-terminal triglycine donor present in large excess to perform bond scission efficiently^32^, otherwise relying on disfavored hydrolytic activity to cleave the bond rather than the enzymatically preferred transpeptidation pathway. These differences explain why the FRET response in A∧*T*-expressing cells shows an inverted response during single protease treatment conditions. Despite these differences in protease efficiency, the expected Boolean-gated FRET responses were achieved.

Here, we have demonstrated that linker topology sets the logical response of a genetically encoded FRET biosensor, yielding YES, OR, and AND gates that compile autonomously from single open reading frames and report in both solution and living cells. Since this responsiveness follows from molecular connectivity rather than input chemistry, the recognition motifs are readily exchangeable. We anticipate that individual gates can be nested into higher-order circuits, extending multi-input sensing to more complex combinations of cellular activity.

## Author Contributions

Ideation and concept were provided by C.A.D., M.L.R. and R.G. Protein expression, protease treatment, and fluorescent intensity data collection for solution-based studies were performed by H.S. and M.K. Mammalian cell culture, transfection, and transposition was completed by J.W.H. Confocal microscopy, flow cytometry, and relevant Python coding was done by J.W.H.

## Notes

C.A.D., M.L.R., and R.G. have filed a patent application (PCT/US2025/031504) related to the work described in this article. The other authors declare no competing interests.

## ACKNOWLEDGEMENTS

Part of this work was conducted with instrumentation provided by the Joint Center for Deployment and Research in Earth Abundant Materials (JCDREAM). Research reported in this publication was generated using the Department of Laboratory Medicine and Pathology Flow Cytometry Core (CC101467) at the University of Washington (UW). The authors gratefully acknowledge Dale Whittington and Josefin Koehn for training and maintenance of the UW Mass Spectrometry Center.

