## Supplementary Information for "Boolean Logic-responsive FRET Biosensors via Genetically Encoded Autonomous Compilation"

### Supplementary Methods

General expression and purification followed the previously reported method[1].

#### Method S1. Protein Expression

Boolean logic gates and protease encoding plasmids were chemically transformed into *Escherichia coli* BL21(DE3). A transformed *E. coli* colony was cultured in lysogeny broth (LB) with kanamycin at 37 ℃ and agitated at 250 revolutions per minute (rpm) until the optical density (OD600) reached 0.5-0.6. Once OD reached 0.5-0.6, Isopropyl β-D-1-thiogalactopyranoside (IPTG) was added into the culture at final concentration 0.5 mM and subsequently cultured at 16 °C and 215 rpm overnight. Cell pellets were collected via centrifugation at 4000xg at 4 ℃ for 20 minutes. Pellets were stored at -80 ℃ until purification.

#### Method S2. Purification of Boolean logic FRET Sensors and Proteases via IMAC and SEC

*Immobilized Metal Affinity Chromatography (IMAC)*
Cell pellets were resuspended in 40mL of Lysis buffer (20nM Tris, 50nM NaCl, 10mM imidazole) and 10uM phenylmethanesulfonyl fluoride(PMSF) on ice. Cells were lysed via sonication on ice (series of 1-second pulse and 2-second rest for 6 minutes at 30% amplitude). The lysed solution was spun down at 4 ℃, 10,000xg, for 45 minutes.

The cell lysate then underwent immobilized metal affinity chromatography (IMAC) on ÄKTA Pure 25L FPLC (Cytiva; Marlborough, MA) with a 5mL HisTrap HP column. In all processes, the flow rate was 5 mL/min. The HisTrap column was equilibrated with 5 column volumes of lysis buffer before sample was bound. The protein-bound column was washed with 8 column volumes of wash buffer (20nM Tris, 50nM NaCl, 15mM imidazole). Proteins were collected by eluting into 8 volumes of elution buffer (20mM Tris, 50mM NaCl, 250mM imidazole; pH 7.5).

Purified linear Boolean logic FRET sensors and proteases were dialyzed in SnakeSkin™ dialysis tubing, 10 kDa molecular weight cut-off (MWCO) (Fischer Scientific; Waltham, MA) against buffer containing 20 mM Tris, 50 mM NaCl, pH = 7.5 for 4 hours and 30 minutes at 4 ℃. The dialysis buffer was exchanged every 1.5 hours. Glycerol was added to the dialyzed products to a final v/v% of 10% and stored at -80 ℃.

*Size Exclusion Chromatography (SEC)*
To eliminate macrocyclic by-products, IMAC-purified cyclic Boolean logic FRET sensors were further purified using a HiLoad 26/600 Superdex 75 pg size-exclusion chromatography (SEC) column on an ÄKTA Pure system. Linear FRET sensors were processed under identical SEC conditions for additional purification. The column was equilibrated with a high-salt buffer (20 mM Tris, 300 mM NaCl, pH 7.5). Samples were loaded at a flow rate of 2.6 mL $min^{-1}$ and subsequently eluted with equilibration buffer at 1.7 mL $min^{-1}$. Eluted fractions were collected in 96-well plates. Fraction purity was assessed by SDS-PAGE prior to pooling (Supplementary Figure 1–3). The pooled protein samples were concentrated using Amicon Ultra-15 centrifugal filters (10 kDa MWCO) and stored at −80 °C in buffer supplemented with 10% glycerol.

**Method S3. In-solution Protease Treatments**

To a buffer containing 1 mM dithiolthreotol (DTT), 1mM $CaCl_2$, 1 mM triglycine, 20 mM tris, and 50 mM NaCl, proteases were added to a final concentration of 2 uM for eSrtA(2A9) and 3 uM for TEV and logic gated proteins were added to a final mass concentration of 1 mg/mL. Solutions were incubated for 4 hours at 37 °C and agitated at 250 rpm.

**Method S4. FRET Index Evaluation**

After protease treatment, fluorescent signals were excited at 479 nm and excitation was collected at 520 nm and 602 nm on plate readers for EGFP($I_{DD}$) and FRET signal($I_{DA}$) respectively. For mScarlet3 ($I_{AA}$), fluorescence was excited at 561 nm and excitation was collected at 602 nm . Relative FRET index was determined as $I_{DA}/(I_{DD}+I_{DA})$ and normalized by non-treated logic gates[2]. Further mass inspection was conducted in SDS-PAGE and Liquid chromatography–mass spectrometry(LC-MS).

**Method S5. Plasmid Cloning via Golden Gate**

Mammalian expression plasmids were inserted into doxycycline inducible Sleeping Beauty destination vectors via Golden Gate Assembly. Linear DNA with properly oriented PaqCI restriction enzyme digest sites were ordered from Twist Bioscience (South San Francisco, CA). PaqCI restriction enzyme and T4 DNA Ligase was purchased from New England Biolabs (Ipswich, MA). Reactions were assembled according to manufacturer protocols. Reactions were run in a thermocycler according to the following protocol: 10 x 37 °C to 16 °C for 10 minutes each, 1x 16 °C for 14 hours, 37 °C for 30 minutes, 60 °C for 5 minutes, hold at 4 °C. Assembly products were transformed into NEB 10 electrocompetent *E. Coli* and cultured on antibiotic-laden LB agar plates overnight at 37 °C. After colonies developed, they were picked into liquid LB culture supplemented with 0.1 mg/mL carbenicillin and cultured overnight. Resulting liquid cultures were miniprepped (Qiagen, Germantown, NJ) and their sequence was verified via whole plasmid sequencing (Azenta, Burlington, MA).

**Method S6. FRET Biosensor Transposition into HEK293T via Sleeping Beauty**

After whole plasmid sequencing verified insert sequence was correct, Human Embryonic Kidney 293T (HEK293T) cells were plated at 40% confluence the day prior to transfection. After allowing them to attach overnight, cells were transfected using Lipofectamine 3000 (ThermoFisher, Waltham, MA) in a 5:1 ratio of biosensor-containing transposon plasmid to SB100X (AddGene #127909). After 24 hours, transfection media was replaced with regular DMEM supplemented with 10% fetal bovine serum (Cytiva, Marlborough, MA) and 1x penicillin/streptomycin (ThermoFisher, Waltham, MA). Two days post-transfection, cells were passaged and plated into DMEM supplemented with a selection antibiotic that corresponds to the transposon they were transfected with (puromycin 10 µg/mL, ThermoFisher Waltham, MA; hygromycin, 100 µg/mL, ThermoFisher Waltham, MA; blasticidin 10 µg/mL, ThermoFisher Waltham, MA). Cells were continuously grown and passaged under antibiotic selection for a period of two weeks post-transfection. Resulting cells were uniformly modified with the biosensors and could be cultured in the long term without antibiotics and without detectable loss in transgene expression.

**Method S7. Protease transfection into HEK293T**

Proteases were expressed via CMV-driven expression after transient transfection with Lipofectamine 3000 (ThermoFisher, Waltham, MA). Plasmids were sourced from Genscript

(Piscataway, NJ) and cloned into pcDNA3.1(+). All proteases were expressed in equimolar quantities with a nuclear-localized emiRFP703 via polycistronic expression with a P2A self-cleaving peptide sequence. For double protease transfections, a single plasmid was cloned with TEV and eSrtA separated by P2A self-cleaving sites upstream of the P2A-emiRFP703. Transfection followed standard protocols from the manufacturer.

On day 0, HEK293T were plated into doxycycline (1 µg/mL, ThermoFisher, Waltham, MA) laden DMEM and allowed to attach overnight. On day 1, cells were transfected with the protease plasmids such that every combination of proteases was represented. For a no protease control, cells were transfected with only emiRFP703. On day 2, the transfected mix was aspirated and replaced with fresh DMEM, and cells were given an additional day to recover. On day 3, cells were taken for further processing and data acquisition as necessary.

**Method S8. Confocal Microscopy of FRET Activity**
On day 0, cells were plated onto gelatin treated glass-bottomed wells (Cellvis, Mountain View, CA) and allowed to attach overnight. Transfections followed previously outlined protocol. On day 3, cells were imaged using a Leica Stellaris (Leica, Wetzlar, DE) such that only the 485 nm laser line was used to excite both EGFP and mScarlet-I3 emission channels. On a separate laser line, a 670 nm laser was used for excitation of emiRFP703. Images were collected in 2048x2048 format using a Z-stack acquisition to remove differences in emission based on working distance from the detector. After collection, images were Z projected to yield a 2D image with maximum intensity values at each pixel.

**Method S9. Image Processing and Analysis**
Z projected images were processed using CellProfiler[3]. Briefly, an image mask was generated using the emiRFP703 channel to isolate the nuclei of cells that were successfully transfected. This mask was applied to both the EGFP and mScarlet3 channels, recorded as previously reported. Relative fret index was calculated by dividing the pixel intensity in the mScarlet3 channel by the sum of the mScarlet3 and EGFP channels and a new image was generated to visualize the relative FRET index at every pixel. Images were then false colored using Viridis color mapping.

**Method S10. Flow Cytometry**
Cells were transfected according to previously outlined protocol. On day 3, cells were dissociated using TrypLE (ThermoFisher, Waltham, MA). After treating with TrypLE for 5 minutes, cells were pipette mixed and added to a suspension of Hanks Buffered Phosphate Solution (HBSS) (ThermoFisher, Waltham, MA) with 1% bovine serum albumin (ThermoFisher, Waltham, MA) and 10 mM HEPES (ThermoFisher, Waltham, MA) in a 1:1 v:v ratio. Cells were pelleted at 200g for 5 minutes and supernatant was aspirated off. Cells were resuspended in HBSS+1%BSA+10mM HEPES and analyzed on a Symphony A3 (BD, Franklin Lakes, NJ).

**Method S11. Python Processing of Flow Cytometry Data**
Code information can be found in the following GitHub repository:
Jack Hoye, University of Washington, “flow_processing”, GitHub, https://github.com/DeForestLab/flow_processing

### Bacterial Construct Amino Acid Sequences Used in this Study

TEV (MBP-6xHis-superTEV-polyR):
MGIEEGKLVIWINGDKGYNGLAEVGKKFEKDTGIKVTVEHPDKLEEKFPQVAATGDGPDIIFWA
HDRFGGYAQSGLLAEITPDKAFQDKLYPFTWDAVRYNGKLIAYPIAVEALSLIYNKDLLPNPPK
TWEEIPALDKELKAKGKSALMFNLQEPYFTWPLIAADGGYAFKYENGKYDIKDVGVDNAGAKAG
LTFLVDLIKNKHMNADTDYSIAEAAFNKGETAMTINGPWAWSNIDTSKVNYGVTVLPTFKGQPS
KPFVGVLSAGINAASPNKELAKEFLENYLLTDEGLEAVNKDKPLGAVALKSYEEELAKDPRIAA
TMENAQKGEIMPNIPQMSAFWYAVRTAVINAASGRQTVDEALKDAQTNSSSNNNNNNNNNNLGI
EGRGDFAGHHHHHHGESLFKGPRDYNPISSTIVHLTNESDGHTTSLYGIGFGPFIITNKHLFR
RNNGTLVVQSLHGVFKVKNTTTLQQHLIDGRDMIIIRMPKDFPPFPQKLKFREPQREERIVLVT
TNFQTKSMSSMVSDTSSTFPSGDGIFWKHWIQTKDGQCGSPLVSTRDGFIVGIHSASNFTNTNN
YFTSVPKNFMELLTNQEAQQWVSGWRLNADSVLWGGHKVFMDKPEEPFQPVKEATQLMNRRRRR
*

eSrtA(2A9) (eSrtA(2A9)-6xHis):
MQAKPQIPKDKSKVAGYIEIPDADIKEPVYPGPATREQLNRGVCFHDENESLDDQNISIAGHTF
IDRPNYQFTNLKAAKPGSMVYFKVGNETRIYKMTSIRKVHPNAVEVLDEQEGKDKQLTLVTCDD
YNEETGVWESRKIFVATEVKGSHHHHHH*

A (EGFP-eSrtA(2A9)c-mScarlet3-6xHis):
MGSSMVSKGEELFTGVVPILVELDGDVNGHKFSVSGEGEGDATYGKLTLKFICTTGKLPVPWPT
LVTTLTYGVQCFSRYPDHMKQHDFFKSAMPEGYVQERTIFFKDDGNYKTRAEVKFEGDTLVNRI
ELKGIDFKEDGNILGHKLEYNYNSHNVYIMADKQKNGIKVNFKIRHNIEDGSVQLADHYQQNTP
IGDGPVLLPDNHYLSTQSALSKDPNEKRDHMVLLEFVTAAGITLGMDELYKGGSGGSGGSGGSG
GSLAETGGGSGGSGGSGGSGGSMDSTEAVIKEFMRFKVHMEGSMNGHEFEIEGEGEGRPYEGTQ
TAKLRVTKGGPLPFSWDILSPQFMYGSRAFTKHPADIPDYWKQSFPEGFKWERVMNFEDGGAVS
VAQDTSLEDGTLIYKVKLRGTNFPPDGPVMQKKTMGWEASTERLYPEDVVLKGDIKMALRLKDG
GRYLADFKTTYRAKKPVQMPGAFNIDRKLDITSHNEDYTVVEQYERSVARHSTGGSGGSGGSGG
SGGSLEHHHHHH*

AvT (EGFP-eSrtA(2A9)c-TEVc-mScarlet3-6xHis):
MGSSMVSKGEELFTGVVPILVELDGDVNGHKFSVSGEGEGDATYGKLTLKFICTTGKLPVPWPT
LVTTLTYGVQCFSRYPDHMKQHDFFKSAMPEGYVQERTIFFKDDGNYKTRAEVKFEGDTLVNRI
ELKGIDFKEDGNILGHKLEYNYNSHNVYIMADKQKNGIKVNFKIRHNIEDGSVQLADHYQQNTP
IGDGPVLLPDNHYLSTQSALSKDPNEKRDHMVLLEFVTAAGITLGMDELYKGGSGGSGGSGGSG
GSLAETGGGSGGSGGSENLYFQSGGSGGSGGSGGSGGSMDSTEAVIKEFMRFKVHMEGSMNGHE
FEIEGEGEGRPYEGTQTAKLRVTKGGPLPFSWDILSPQFMYGSRAFTKHPADIPDYWKQSFPEG
FKWERVMNFEDGGAVSVAQDTSLEDGTLIYKVKLRGTNFPPDGPVMQKKTMGWEASTERLYPED
VVLKGDIKMALRLKDGGRYLADFKTTYRAKKPVQMPGAFNIDRKLDITSHNEDYTVVEQYERSV
ARHSTGGSGGSGGSGGSGGSLEHHHHHH*

AΛT (CfaC-Ext.C-EGFP-eSrtA(2A9)c-mScarlet3-TEVc-6xHis-Ext.N-CfaN-SsrA):
MGSSVKIISRKSLGTQNVYDIGVEKDHNFLLKNGLVASNCFNGGSGGSGGSGGSGGSMVSKGEE
LFTGVVPILVELDGDVNGHKFSVSGEGEGDATYGKLTLKFICTTGKLPVPWPTLVTTLTYGVQC
FSRYPDHMKQHDFFKSAMPEGYVQERTIFFKDDGNYKTRAEVKFEGDTLVNRIELKGIDFKEDG
NILGHKLEYNYNSHNVYIMADKQKNGIKVNFKIRHNIEDGSVQLADHYQQNTPIGDGPVLLPDN

HYLSTQSALSKDPNEKRDHMVLLEFVTAAGITLGMDELYKGGSGGSGGSGGSGGSLAETGGGSG
GSGGSGGSGGSMDSTEAVIKEFMRFKVHMEGSMNGHEFEIEGEGEGRPYEGTQTAKLRVTKGGP
LPFSWDILSPQFMYGSRAFTKHPADIPDYWKQSFPEGFKWERVMNFEDGGAVSVAQDTSLEDGT
LIYKVKLRGTNFPPDGPVMQKKTMGWEASTERLYPEDVVLKGDIKMALRLKDGGRYLADFKTTY
RAKKPVQMPGAFNIDRKLDITSHNEDYTVVEQYERSVARHSTGGSGGSGGSGGSGGSGGSGGSE
NLYFQSGGSGGSGGSGGSGGSHHHHHHGGSGGSGGSGGSGGSAEYCLSYDTEILTVEYGFLPIG
KIVEERIECTVYTVDKNGFVYTQPIAQWHNRGEQEVFEYCLEDGSIIRATKDHKFMTTDGQMLP
IDEIFERGLDLKQVDGLPAANDENYALAA*

A (EGFP-eSrtA(2A9)c-mScarlet3-6xHis):

MGSSMVSKGEELFTGVVPILVELDGDVNGHKFSVSGEGEGDATYGKLTLKFICTTGKLPVPWPT
LVTTLTYGVQCFSRYPDHMKQHDFFKSAMPEGYVQERTIFFKDDGNYKTRAEVKFEGDTLVNRI
ELKGIDFKEDGNILGHKLEYNYNSHNVYIMADKQKNGIKVNFKIRHNIEDGSVQLADHYQQNTP
IGDGPVLLPDNHYLSTQSALSKDPNEKRDHMVLLEFVTAAGITLGMDELYKGGSGGSGGSGGSG
GSLAETGGGSGGSGGSGGSGGSMDSTEAVIKEFMRFKVHMEGSMNGHEFEIEGEGEGRPYEGTQ
TAKLRVTKGGPLPFSWDILSPQFMYGSRAFTKHPADIPDYWKQSFPEGFKWERVMNFEDGGAVS
VAQDTSLEDGTLIYKVKLRGTNFPPDGPVMQKKTMGWEASTERLYPEDVVLKGDIKMALRLKDG
GRYLADFKTTYRAKKPVQMPGAFNIDRKLDITSHNEDYTVVEQYERSVARHSTGGSGGSGGSGG
SGGSLEHHHHHH*

A∨T (EGFP-eSrtA(2A9)c-TEVc-mScarlet3-6xHis):

MGSSMVSKGEELFTGVVPILVELDGDVNGHKFSVSGEGEGDATYGKLTLKFICTTGKLPVPWPT
LVTTLTYGVQCFSRYPDHMKQHDFFKSAMPEGYVQERTIFFKDDGNYKTRAEVKFEGDTLVNRI
ELKGIDFKEDGNILGHKLEYNYNSHNVYIMADKQKNGIKVNFKIRHNIEDGSVQLADHYQQNTP
IGDGPVLLPDNHYLSTQSALSKDPNEKRDHMVLLEFVTAAGITLGMDELYKGGSGGSGGSGGSG
GSLAETGGGSGGSGGSENLYFQSGGSGGSGGSGGSGGSMDSTEAVIKEFMRFKVHMEGSMNGHE
FEIEGEGEGRPYEGTQTAKLRVTKGGPLPFSWDILSPQFMYGSRAFTKHPADIPDYWKQSFPEG
FKWERVMNFEDGGAVSVAQDTSLEDGTLIYKVKLRGTNFPPDGPVMQKKTMGWEASTERLYPED
VVLKGDIKMALRLKDGGRYLADFKTTYRAKKPVQMPGAFNIDRKLDITSHNEDYTVVEQYERSV
ARHSTGGSGGSGGSGGSGGSLEHHHHHH*

A∧T (CfaC-Ext.C-EGFP-eSrtA(2A9)c-mScarlet3-TEVc-6xHis-Ext.N-CfaN-SsrA):

MGSSVKIISRKSLGTQNVYDIGVEKDHNFLLKNGLVASNCFNGGSGGSGGSGGSGGSMVSKGEE
LFTGVVPILVELDGDVNGHKFSVSGEGEGDATYGKLTLKFICTTGKLPVPWPTLVTTLTYGVQC
FSRYPDHMKQHDFFKSAMPEGYVQERTIFFKDDGNYKTRAEVKFEGDTLVNRIELKGIDFKEDG
NILGHKLEYNYNSHNVYIMADKQKNGIKVNFKIRHNIEDGSVQLADHYQQNTPIGDGPVLLPDN
HYLSTQSALSKDPNEKRDHMVLLEFVTAAGITLGMDELYKGGSGGSGGSGGSGGSLAETGGGSG
GSGGSGGSGGSMDSTEAVIKEFMRFKVHMEGSMNGHEFEIEGEGEGRPYEGTQTAKLRVTKGGP
LPFSWDILSPQFMYGSRAFTKHPADIPDYWKQSFPEGFKWERVMNFEDGGAVSVAQDTSLEDGT
LIYKVKLRGTNFPPDGPVMQKKTMGWEASTERLYPEDVVLKGDIKMALRLKDGGRYLADFKTTY
RAKKPVQMPGAFNIDRKLDITSHNEDYTVVEQYERSVARHSTGGSGGSGGSGGSGGSGGSGGSE
NLYFQSGGSGGSGGSGGSGGSHHHHHHGGSGGSGGSGGSGGSAEYCLSYDTEILTVEYGFLPIG
KIVEERIECTVYTVDKNGFVYTQPIAQWHNRGEQEVFEYCLEDGSIIRATKDHKFMTTDGQMLP
IDEIFERGLDLKQVDGLPAANDENYALAA*

**Mammalian Construct Amino Acid Sequences Used in this Study**

A (EGFP-eSrtA(2A9)c-mScarlet3-6xHis):
MGSSMVSKGEELFTGVVPILVELDGDVNGHKFSVSGEGEGDATYGKLTLKFICTTGKLPVPWPT
LVTTLTYGVQCFSRYPDHMKQHDFFKSAMPEGYVQERTIFFKDDGNYKTRAEVKFEGDTLVNRI
ELKGIDFKEDGNILGHKLEYNYNSHNVYIMADKQKNGIKVNFKIRHNIEDGSVQLADHYQQNTP
IGDGPVLLPDNHYLSTQSALSKDPNEKRDHMVLLEFVTAAGITLGMDELYKGGSGGSGGSGGSG
GSLAETGGGSGGSGGSGGSGGSMDSTEAVIKEFMRFKVHMEGSMNGHEFEIEGEGEGRPYEGTQ
TAKLRVTKGGPLPFSWDILSPQFMYGSRAFTKHPADIPDYWKQSFPEGFKWERVMNFEDGGAVS
VAQDTSLEDGTLIYKVKLRGTNFPPDGPVMQKKTMGWEASTERLYPEDVVLKGDIKMALRLKDG
GRYLADFKTTYRAKKPVQMPGAFNIDRKLDITSHNEDYTVVEQYERSVARHSTGGSGGSGGSGG
SGGSLEHHHHHH*

A∨T (EGFP-eSrtA(2A9)c-TEVc-mScarlet3-6xHis):
MGSSMVSKGEELFTGVVPILVELDGDVNGHKFSVSGEGEGDATYGKLTLKFICTTGKLPVPWPT
LVTTLTYGVQCFSRYPDHMKQHDFFKSAMPEGYVQERTIFFKDDGNYKTRAEVKFEGDTLVNRI
ELKGIDFKEDGNILGHKLEYNYNSHNVYIMADKQKNGIKVNFKIRHNIEDGSVQLADHYQQNTP
IGDGPVLLPDNHYLSTQSALSKDPNEKRDHMVLLEFVTAAGITLGMDELYKGGSGGSGGSGGSG
GSLAETGGGSGGSGGSENLYFQSGGSGGSGGSGGSGGSMDSTEAVIKEFMRFKVHMEGSMNGHE
FEIEGEGEGRPYEGTQTAKLRVTKGGPLPFSWDILSPQFMYGSRAFTKHPADIPDYWKQSFPEG
FKWERVMNFEDGGAVSVAQDTSLEDGTLIYKVKLRGTNFPPDGPVMQKKTMGWEASTERLYPED
VVLKGDIKMALRLKDGGRYLADFKTTYRAKKPVQMPGAFNIDRKLDITSHNEDYTVVEQYERSV
ARHSTGGSGGSGGSGGSGGSLEHHHHHH*

A∧T (CfaC-EGFP-eSrtA(2A9)c-mScarlet3-TEVc-CfaN):
MGSSVKIISRKSLGTQNVYDIGVEKDHNFLLKNGLVASNCFNGGSGGSGGSGGSGGSMVSKGEE
LFTGVVPILVELDGDVNGHKFSVSGEGEGDATYGKLTLKFICTTGKLPVPWPTLVTTLTYGVQC
FSRYPDHMKQHDFFKSAMPEGYVQERTIFFKDDGNYKTRAEVKFEGDTLVNRIELKGIDFKEDG
NILGHKLEYNYNSHNVYIMADKQKNGIKVNFKIRHNIEDGSVQLADHYQQNTPIGDGPVLLPDN
HYLSTQSALSKDPNEKRDHMVLLEFVTAAGITLGMDELYKGGSGGSGGSGGSGGSLAETGGGSG
GSGGSGGSGGSMDSTEAVIKEFMRFKVHMEGSMNGHEFEIEGEGEGRPYEGTQTAKLRVTKGGP
LPFSWDILSPQFMYGSRAFTKHPADIPDYWKQSFPEGFKWERVMNFEDGGAVSVAQDTSLEDGT
LIYKVKLRGTNFPPDGPVMQKKTMGWEASTERLYPEDVVLKGDIKMALRLKDGGRYLADFKTTY
RAKKPVQMPGAFNIDRKLDITSHNEDYTVVEQYERSVARHSTGGSGGSGGSGGSGGSGGSGGSE
NLYFQSGGSGGSGGSGGSGGSHHHHHHGGSGGSGGSGGSGGSAEYCLSYDTEILTVEYGFLPIG
KIVEERIECTVYTVDKNGFVYTQPIAQWHNRGEQEVFEYCLEDGSIIRATKDHKFMTTDGQMLP
IDEIFERGLDLKQVDGLPAANDENYALAA*

TEV Protease (TEV-NLS-P2A-emiRFP703-NLS)
MGSSGESLFKGPRDYNPISSTIVHLTNESDGHTTSLYGIGFGPFIITNKHLFRRNNGTLVVQSL
HGVFKVKNTTTLQQHLIDGRDMIIIRMPKDFPPFPQKLKFREPQREERIVLVTTNFQTKSMSSM
VSDTSSTFPSGDGIFWKHWIQTKDGQCGSPLVSTRDGFIVGIHSASNFTNTNNYFTSVPKNFME
LLTNQEAQQWVSGWRLNADSVLWGGHKVFMDKPEEPFQPVKEATQLMNRRRRRGSGSATNFSLL
KQAGDVEENPGPGSGSKRPAATKKAGQAKKKKAEGSVARQPDLLTCEHEEIHLAGSIQPHGALL
VVSEHDHRVIQASANAAEFLNLGSVLGVPLAEIDGDLLIKILPHLDPTAEGMPVAVRCRIGNPS
TEYCGLMHRPPEGGLIIELERAGPSIDLSGTLAPALERIRTAGSLRALCDDTVLLFQQCTGYDR
VMVYRFDEQGHGLVFSECHVPGLESYFGNRYPSSLVPQMARQLYVRQRVRVLDVTYQPVPLEP

RLSPLTGRDLDMSGCFLRSMSPIHLQFLKDMGVRATLAVSLVVGGKLWGLVVCHHYLPRFIRFE
LRAICKRLAERIATRITALESPKKKRKV*

eSrtA(2A9) Protease (eSrtA-NLS-P2A-emiRFP703-NLS)
MGSSMQAKPQIPKDKSKVAGYIEIPDADIKEPVYPGPATREQLNRGVCFHDENESLDDQNISIA
GHTFIDRPNYQFTNLKAAKPGSMVYFKVGNETRIYKMTSIRKVHPNAVEVLDEQEGKDKQLTLV
TCDDYNEETGVWESRKIFVATEVKGSGSATNFSLLKQAGDVEENPGPGSGSKRPAATKKAGQAK
KKKAEGSVARQPDLLTCEHEEIHLAGSIQPHGALLVVSEHDHRVIQASANAAEFLNLGSVLGVP
LAEIDGDLLIKILPHLDPTAEGMPVAVRCRIGNPSTEYCGLMHRPPEGGLIIELERAGPSIDLS
GTLAPALERIRTAGSLRALCDDTVLLFQQCTGYDRVMVYRFDEQGHGLVFSECHVPGLESYFGN
RYPSSLVPQMARQLYVRQRVRVLVDVTYQPVPLEPRLSPLTGRDLDMSGCFLRSMSPIHLQFLK
DMGVRATLAVSLVVGGKLWGLVVCHHYLPRFIRFELRAICKRLAERIATRITALESPKKKRKV*

TEV + eSrtA(2A9) Proteases (TEV-P2A-eSrtA-NLS-P2A-emiRFP703-NLS)
MGSSGESLFKGPRDYNPISSTIVHLTNESDGHTTSLYGIGFGPFIITNKHLFRRNNGTLVVQSL
HGVFKVKNTTTLQQHLIDGRDMIIIRMPKDFPPFPQKLKFREPQREERIVLVTTNFQTKSMSSM
VSDTSSTFPSGDGIFWKHWIQTKDGQCGSPLVSTRDGFIVGIHSASNFTNTNNYFTSVPKNFME
LLTNQEAQQWVSGWRLNADSVLWGGHKVFMDKPEEPFQPVKEATQLMNRRRRRGSGSATNFSLL
KQAGDVEENPGPGSGSMQAKPQIPKDKSKVAGYIEIPDADIKEPVYPGPATREQLNRGVCFHDE
NESLDDQNISIAGHTFIDRPNYQFTNLKAAKPGSMVYFKVGNETRIYKMTSIRKVHPNAVEVLD
EQEGKDKQLTLVTCDDYNEETGVWESRKIFVATEVKGSGSGSATNFSLLKQAGDVEENPGPGSG
SKRPAATKKAGQAKKKKAEGSVARQPDLLTCEHEEIHLAGSIQPHGALLVVSEHDHRVIQASAN
AAEFLNLGSVLGVPLAEIDGDLLIKILPHLDPTAEGMPVAVRCRIGNPSTEYCGLMHRPPEGGL
IIELERAGPSIDLSGTLAPALERIRTAGSLRALCDDTVLLFQQCTGYDRVMVYRFDEQGHGLVF
SECHVPGLESYFGNRYPSSLVPQMARQLYVRQRVRVLVDVTYQPVPLEPRLSPLTGRDLDMSGC
FLRSMSPIHLQFLKDMGVRATLAVSLVVGGKLWGLVVCHHYLPRFIRFELRAICKRLAERIATR
ITALESPKKKRKV*

**Figure S1 SEC Analysis of A Biosensor**

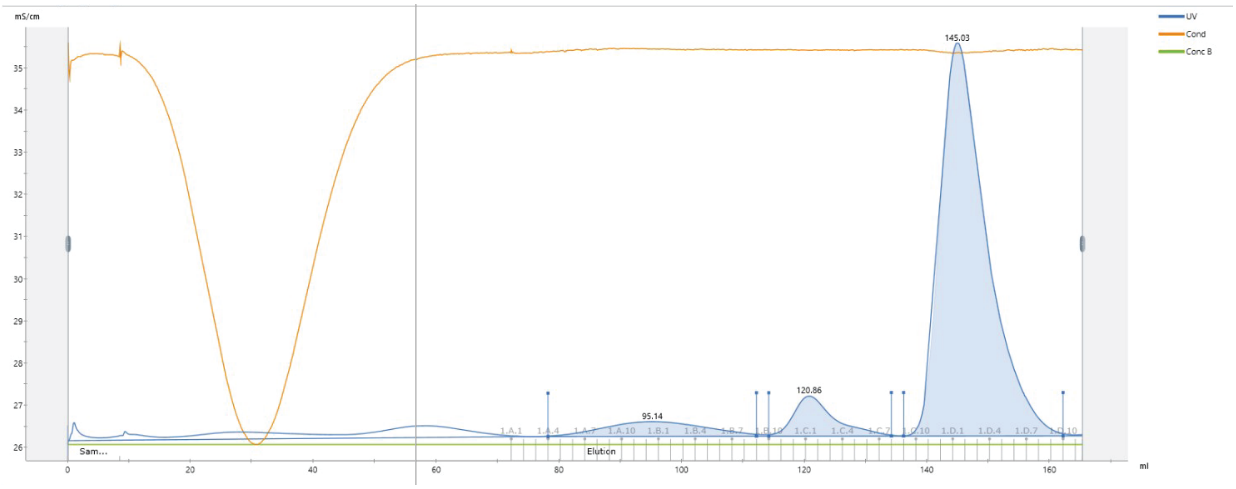

SEC chromatogram for the A biosensor. Fractions were collected for subsequent analysis based on UV absorbance.

**Figure S2 SDS-PAGE Analysis of Purified A Biosensor**

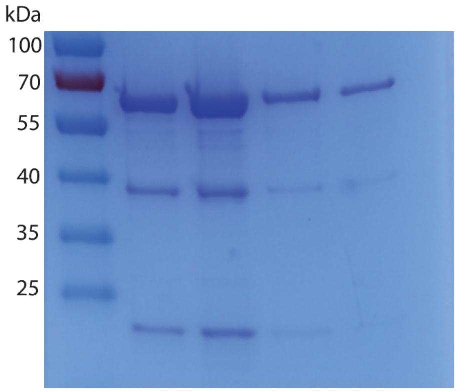

SDS-PAGE analysis of purified A biosensor sample. Each lane corresponds to a different fraction in the same dominant peak collected in SEC (~145 mL). (Expected Mass: 57.1kDa)

**Figure S3 LC-MS Analysis of Purified A Biosensor**

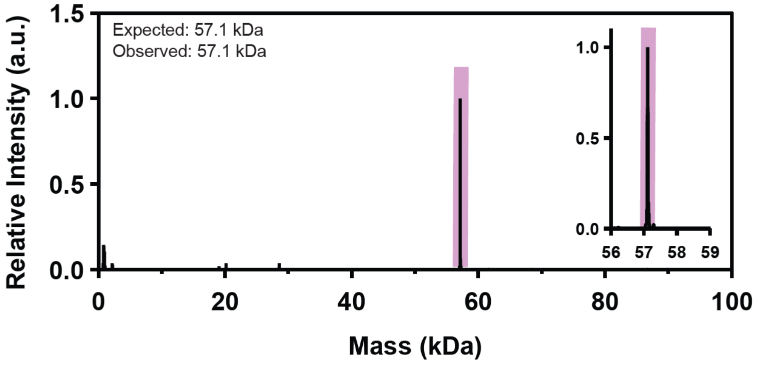

LC-MS analysis of SEC-purified A biosensor. Expected and observed masses are listed on the top-left of the plot. Inset graphs depict a zoomed-in view of the LC-MS peak.

**Figure S4 SEC Analysis of AVT Biosensor**

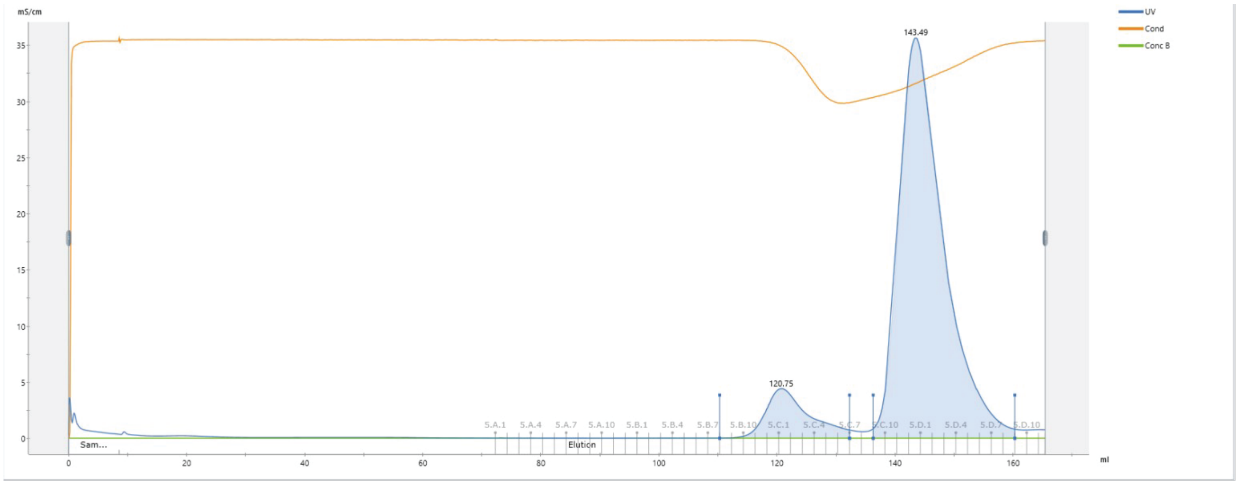

SEC chromatogram for the AVT biosensor. Fractions were collected for subsequent analysis based on UV absorbance.

**Figure S5 SDS-PAGE Analysis of Purified AᴠT Biosensor**

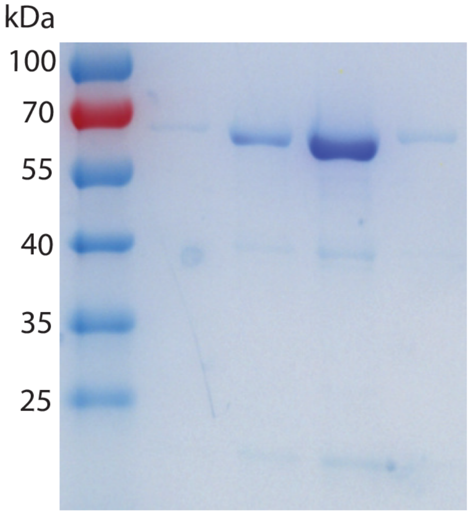

SDS-PAGE gel showing purified AᴠT biosensor. Each lane corresponds to a different fraction in the same dominant peak collected in SEC (~141 mL). (Expected Mass: 58.6kDa)

**Figure S6 LC-MS Analysis of Purified AVT Biosensor**

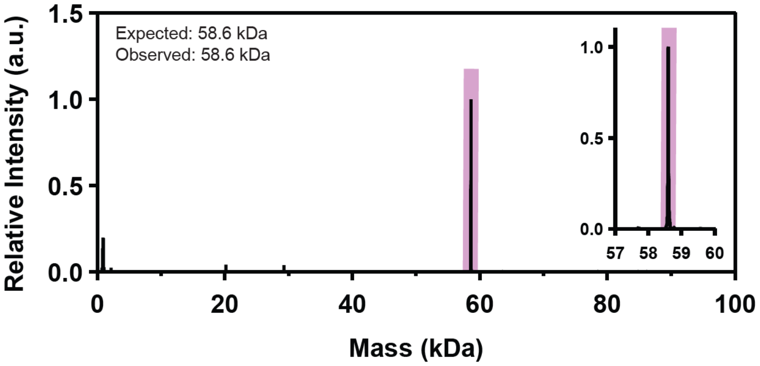

LC-MS analysis of SEC-purified AVT biosensor. Expected and observed masses are listed on the top-left of the plot. Inset graphs depict a zoomed-in view of the LC-MS peak.

**Figure S7 SEC Analysis of A∧T Biosensor**

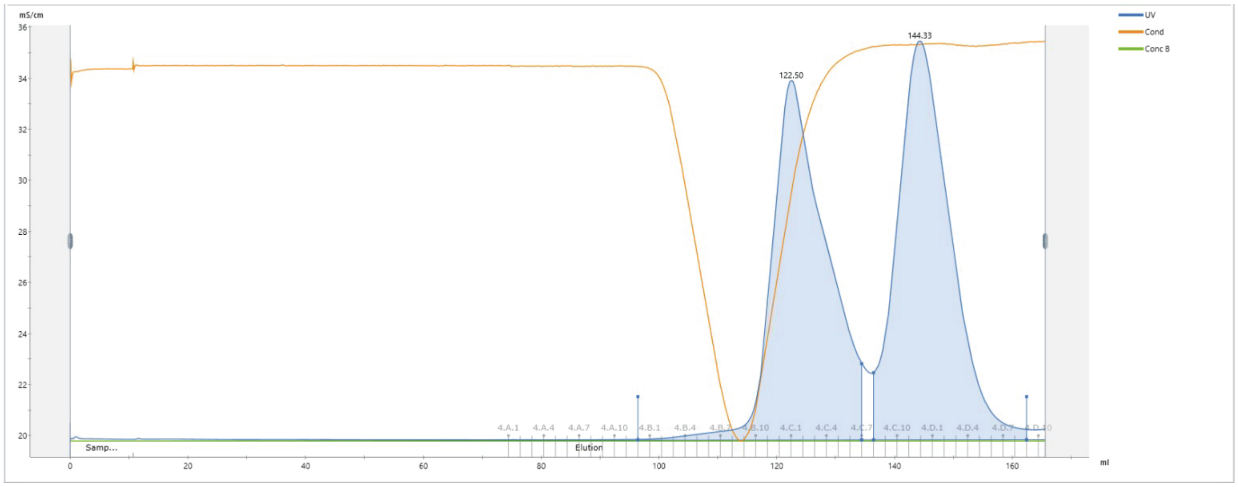

SEC chromatogram for the A∧T biosensor. Fractions were collected for subsequent analysis based on UV absorbance.

**Figure S8 SDS-PAGE Analysis of Purified AΛT Biosensor**

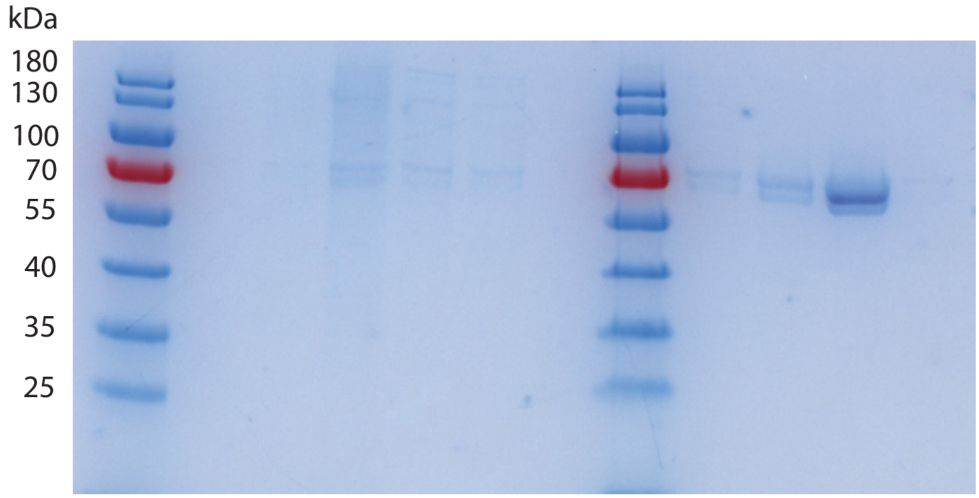

SDS-PAGE analysis of purified AΛT biosensor sample. The first eluting SEC peak is analyzed on the left; the second on the right. Each lane corresponds to a different fraction in the same dominant peak collected in SEC (left ~122 mL, right ~144 mL). Fractions from the second peak were selected for future use. (Expected Mass: 61.7kDa)

**Figure S9 LC-MS Analysis of Purified A∧T Biosensor**

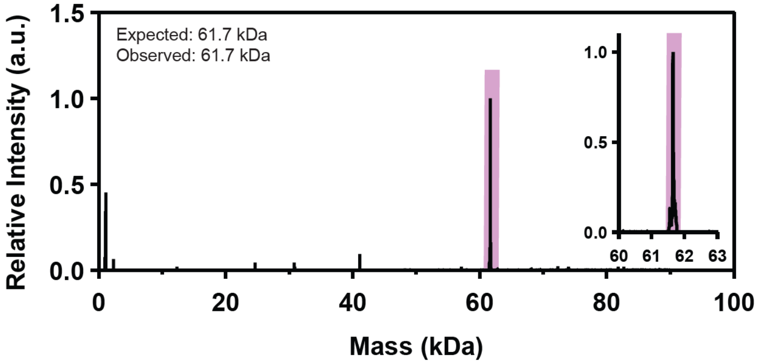

LC-MS analysis of SEC-purified A∧T biosensor. Expected and observed masses are listed on the top-left of the plot. Inset graphs depict a zoomed-in view of the LC-MS peak.

### Figure S10 Flow Cytometry Gating Strategy

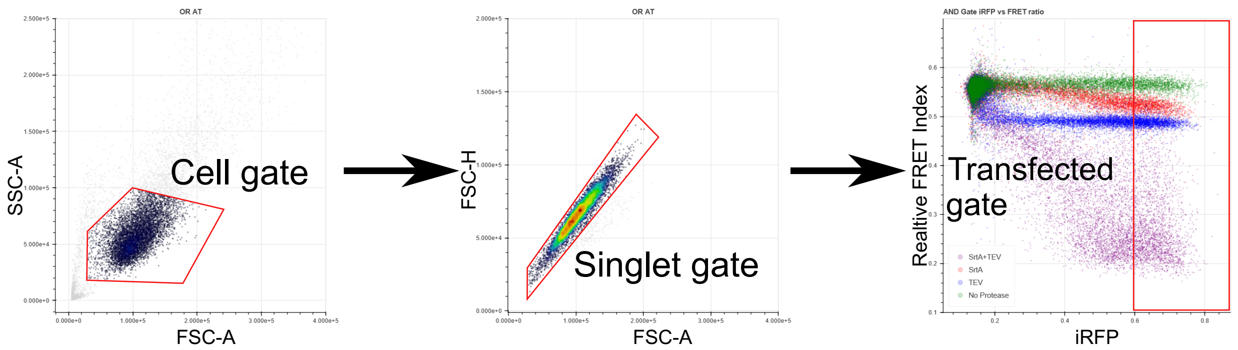

Cells are dissociated using TrypLE and cell strained (50 µm) to remove large clusters. After this, cells are separated from large clusters and cell debris using the FSC-A vs. SSC-A gate as shown. Daughter to this gate is the singlet gate, shown on the FSC-A vs. FSC-H, which further isolates single cells against small cell clusters. Both of these gates allowed ~90% of events to pass through. FRET ratio was then calculated using FlowCal package in Python, which allows for an event-by-event calculation of the FRET ratio according to previously mentioned relative FRET index calculations. Calculated FRET index was plotted against the iRFP channel to identify what expression level of transfected proteases is sufficient to enable complete gate scission. This iRFP gate is then added to the gating strategy in FlowKit python package and gated data is exported as a .CSV for further analysis.

**Figure S11 Confocal Image Processing**

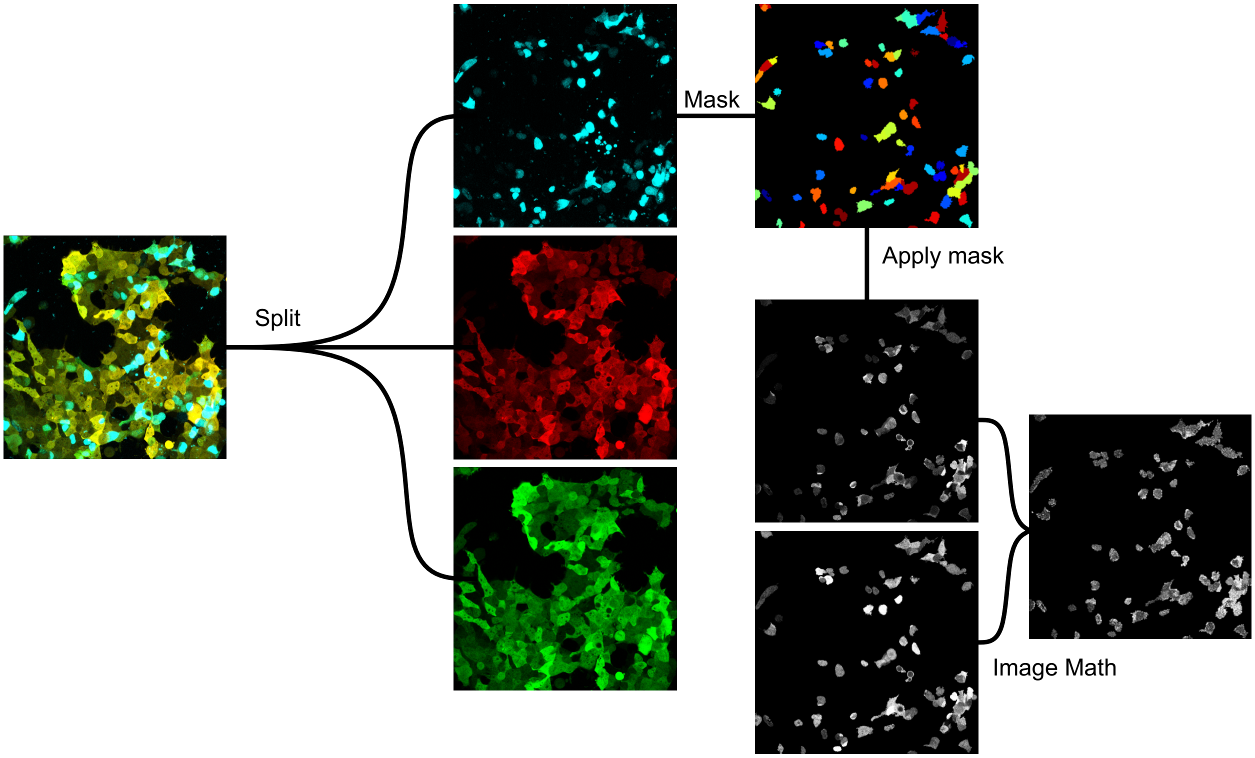

To eliminate background and isolate cells in the population that have been transfected, the first step of the CellProfiler pipeline is to identify primary objects in the iRFP channel (cyan). Next, these objects are used as a mask over the EGFP (green) and mScarlet-3 (red) channels, with pixels outside of the identified objects having their intensity set to 0. Image math operations first calculate the FRET index denominator ($I_{red}+I_{green}$, i.e., the intensity of the acceptor and donor fluorescence, respectively) in a pixel wise manner. Next, this denominator is used on the red channel, yielding the relative FRET index at each relevant pixel. Image is then exported from CellProfiler as a .PNG and false colored using ImageJ.
